# CryoNeRF: reconstruction of homogeneous and heterogeneous cryo-EM structures using neural radiance field

**DOI:** 10.1101/2025.01.10.632460

**Authors:** Huaizhi Qu, Xiao Wang, Yuanyuan Zhang, Sheng Wang, William Stafford Noble, Tianlong Chen

**Author notes:** **Materials & Correspondence** Sheng Wang, William Stafford Noble, Tianlong Chen. These authors contributed equally (Huaizhi Qu, Xiao Wang).

## Abstract

Cryogenic electron microscopy (cryo-EM) has become a widely used technique for determining the 3D structures of proteins. However, cryo-EM datasets often exhibit heterogeneity, with protein particle images from multiple conformations or compositional states. Here we propose **CryoNeRF**, a novel neural radiance fields (NeRF)-based cryo-EM reconstruction framework operating directly in Euclidean 3D space. CryoNeRF introduces a multi-resolution hash encoding and heterogeneity-aware cryo-EM encoder to model cryo-EM heterogenity. Extensive experiments demonstrate the stability and superior performance of CryoNeRF in both homogeneous and heterogeneous settings. On homogeneous datasets, CryoNeRF achieves 15.8% improvement over previous state-of-the-art methods. On both simulated and experimental heterogeneous datasets, CryoNeRF demonstrates exceptional capability in handling both conformational and compositional variations, which is consistent with previous experimental discoveries. Notably, CryoNeRF successfully distinguishes assembly states that even only account for 2% particles of the dataset in cases of compositional heterogeneity.

## Main

Proteins are dynamic macromolecules, and these dynamics are directly linked to protein function^1-3^. The dynamic nature of a particular protein species will lead to structural heterogeneity within the population observed in a sample because individual protein particles may be in different states when imaging and adopt different 3D conformations. To gain a comprehensive understanding of how proteins function in biological processes, it is crucial to determine their various 3D structures throughout their functional cycles^4-10^.

In the last decade, cryogenic electron microscopy (cryo-EM) has become one of the most popular technologies for 3D structure determination because of its ability to characterize large macromolecules^11-15^; however, resolving heterogeneity of structures in the protein population during one cryo-EM observation remains challenging. In the cryo-EM protocol, samples are frozen in a thin film of vitreous ice, fixing the structures of individual proteins for imaging. Subsequently, transmission electron microscopy (TEM) captures hundreds of images of the film, each image containing numerous particles of a specific type of protein. Captured particles within the dataset may adopt either a common (homogeneous) 3D structure or multiple distinct (heterogeneous) 3D structures, allowing for the assessment of the protein’s structural heterogeneity. Protein heterogeneity can be categorized into two distinct types: conformational heterogeneity, where protein particles of a single protein class undergo continuous motion during capture, and compositional heterogeneity, where protein particles represent different protein classes that have distinct structures. Cryo-EM reconstruction aims to infer the 3D density of protein particles in a dataset from a collection of 2D particle images. Traditional 3D reconstruction methods require expert experience and knowledge to estimate the movement of the target protein. Multibody refinement^16^, implemented in RELION^17^, models protein dynamics by manually selecting independently moving regions on the protein and dividing them into moving rigid bodies. 3D Variability Analysis (3DVA)^18^, implemented in the cryoSPARC^19^ software, models the 3D structures of the protein as a linear combination of a 3D base structure plus a weighted sum of 3D basis functions. Both methods require manual adjustments for better 3D reconstruction, such as manually selected moving regions in multibody refinement and the user-defined number of 3D basis functions in 3DVA. These adjustments greatly increase reconstruction time and require expert knowledge.

Recently, deep neural networks (DNNs) have become widely used to automatically model complex distributions for better 3D reconstruction. For example, a recent method, cryoDRGN^20^, uses a DNN-based autoencoder to model the heterogeneity of protein structures. DNNs are known for their capacity to project complicated relationships into latent embeddings, which allows cryoDRGN to learn the underlying heterogeneity of 3D structures within the population of a cryo-EM dataset. Another DNN-based method, e2gmm^21^, utilizes a DNN to predict the 3D coordinates, amplitude, and width of a Gaussian mixture model (GMM) to represent the density of a protein. However, there still exist several major limitations in existing DNN methods. For instance, the Fourier transformation in cryoDRGN and e2gmm will lead to numerical error and thus information loss of high-frequency features. Additionally, e2gmm adopts a two-stage training design, which makes the training process less efficient and less stable.

To resolve these limitations, we propose CryoNeRF, a DNN cryo-EM reconstruction method that operates in Euclidean 3D space with no Fourier transformation. Neural radiance field (NeRF) is a deep learning method that reconstructs a 3D representation of an object from 2D images captured at different angles. We hypothesized that NeRF could be used in cryo-EM analysis to reconstruct the 3D density of protein particles from 2D projections. In addition, to handle the heterogeneity, we proposed novel multi-resolution hash encoding and heterogeneity-aware cryo-EM encoders. CryoNeRF takes as input 2D particle images derived from EM maps. These images are processed by our heterogeneity-aware cryo-EM encoder to generate a set of particle embeddings of the images. These embeddings, along with the query coordinates in multi-resolution hash encodings, are then fed into NeRF to generate density predictions of the query locations. Using rotation angles and translations of particles in the input images inferred by *ab initio* reconstruction, CryoNeRF can be optimized in an unsupervised fashion by backpropagation of the reconstruction loss between the input images and projections of the predicted 3D maps.

Through extensive benchmark, we observed the proposed method yields stable and outstanding performance in both homogeneous and heterogeneous settings. First, on homogeneous reconstruction dataset, CryoNeRF achieves significant improvement over previous state-of-the-art methods (0.9 Å). On simulated heterogenity setting, CryoNeRF could accurately capture the rotational continuum in the embedding space for conformational heterogenity, and capture the structural similarities and differences of different protein classes for compositional heterogenity. On experimental heterogenity setting, CryoNeRF could accurately map dynamic conformational transitions into a well-organized latent space for conformational heterogeneity, identify distinct assembly states for compositional heterogenity, both yield high consistency with experimental results^22,23.^ Overall, the novel design of CryoNeRF greatly enhances our ability to study biomolecular structures, providing a stable framework for uncovering dynamic processes and assembly mechanisms.

## Results

### Overview of CryoNeRF

CryoNeRF reconstructs 3D density in Euclidean 3D space by using 2D particle images as input. This idea was initially inspired by the neural radiance field (NeRF)^24^, which can leverage projections of a single density from multiple view angles (**Fig. 1a**) to reconstruct the underlying 3D particle. Specifically, CryoNeRF addresses two distinct scenarios in cryo-EM (**Fig. 1b**): homogeneous reconstruction, where the particles captured in the images share an identical class (the same spatial shape) but differ in rotation and translation; and heterogeneous reconstruction, where particles are in different classes (different spatial shapes) as well as in rotation and translation.

**Fig. 1.**
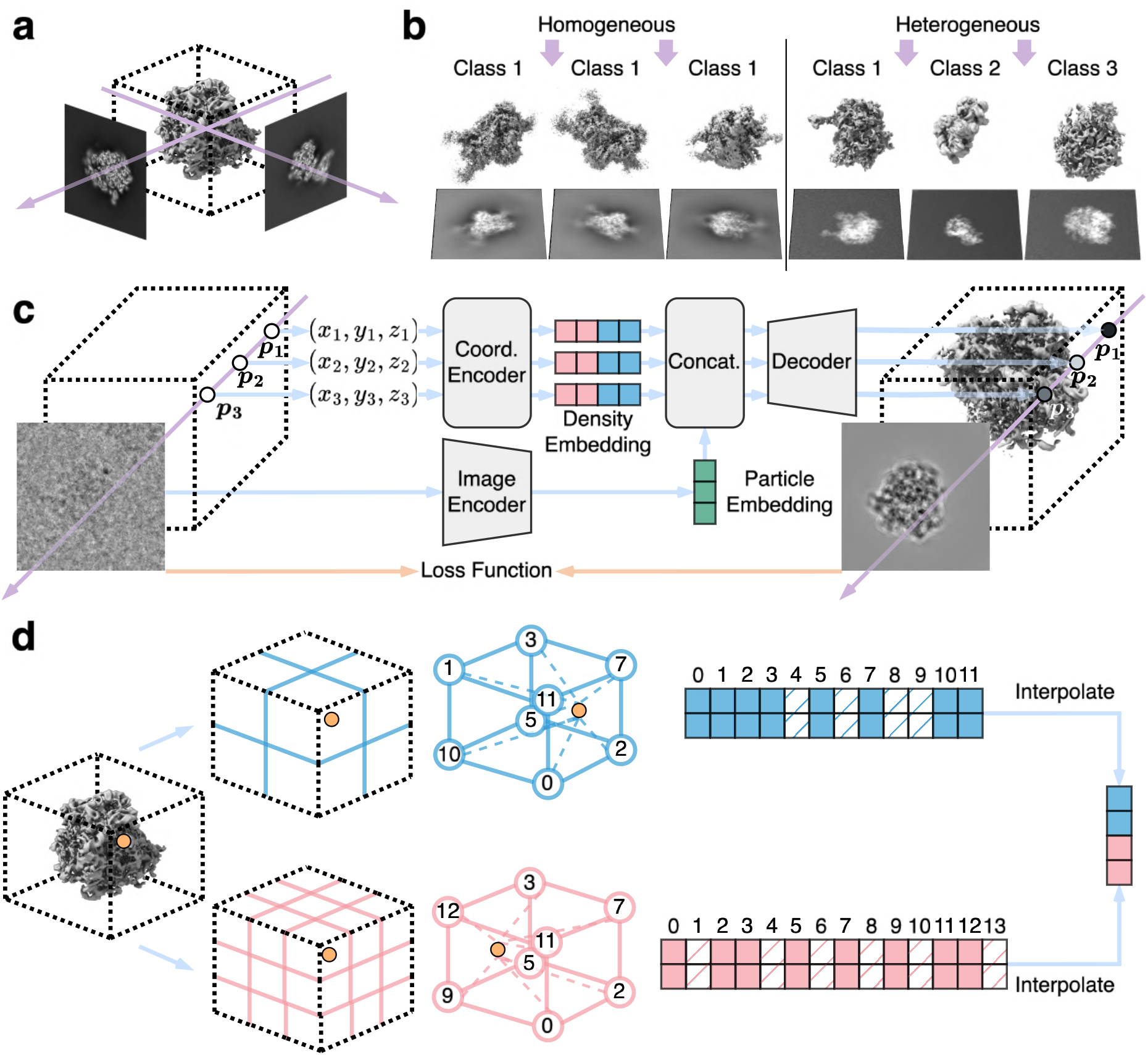
Overview of the CryoNeRF pipeline. **a**, Projections of a protein density at different view angles. **b**, Comparison between homogeneous and heterogeneous cryo-EM datasets. A cryo-EM dataset consists of many 2D projections of 3D protein particles, with each 2D image capturing a 3D particle. In a homogeneous dataset, all the protein particles are from the same class; thus, the available 2D projections are views of the same protein class at different angles. In a heterogeneous dataset, protein particles come from different classes, and the 2D projections are views of many classes of proteins at different angles. **c**, Cryo-EM reconstruction in Euclidean 3D space. The coordinates of points are input into an embedding query module to predict the density embeddings. In parallel, an image encoder extracts particle embeddings from the particle image. A decoder takes both the density and the particle embeddings to predict the density values at each point. After predicting all points, the density is projected and compared with the particle image to compute the loss and optimize the model. **d**, Multi-resolution hash encoding for embedding query. Given a query point, its coordinates are used to look up voxels that contain the point from different spatial resolutions. The vertices of a voxel are hashed to query embeddings from the hash table corresponding to that resolution. Then the embeddings from the voxel vertices are interpolated to obtain the embedding of the query point at the given spatial resolution.

The overall framework of CryoNeRF is illustrated in **Fig. 1c**. Neural radiance field (NeRF), a 3D computer vision technique that operates in Euclidean 3D space and learns the spatial distribution of the field, naturally aligns with cryo-EM reconstruction, which aims to reconstruct the density of proteins. CryoNeRF takes the projected 2D image from cryo-EM and the coordinate of a query point as the input. Here the query point could be any point in the 3D space to query the corresponding density. Using multi-resolution hash encoding (**Fig. 1d**), the model generates the density embedding of the query point. In parallel, the particle embedding of the projected 2D image is extracted by an image encoder. Then the decoder takes the combined embeddings as the input and finally outputs the density of the query points. The overall framework is then optimized by comparing the projection of the reconstructed density with the 2D particle image (see Methods for more details).

To effectively query density embeddings from input coordinates, we use multi-resolution hash encoding^25^. In this technique, the space of interest is represented by several levels of voxel-based representations that divide the space into different granularities of voxels (**Fig. 1d**), where each level divides the space into voxels of different sizes corresponding to different spatial resolutions. Each level is associated with a hash table that stores learnable embedding vectors for its voxel representations.

The density value at a given point is predicted using its coordinates as a query to the multi-resolution hash encoding, retrieving an embedding vector by interpolation of embedding vectors from nearby vertices of the corresponding voxel. This embedding vector is then combined with other information to predict the point’s density (**Fig.1c**).

The voxel representation, by dividing the space into many non-overlapping regions, effectively captures the intricate details of protein density within each voxel and therefore enables high-resolution and precise modeling of heterogeneity. As the hash encoding at each level of resolution enables rapid lookup of the density embedding for a voxel, it enhances the efficiency of querying density embeddings from voxel representations using coordinates.

CryoNeRF is trained end-to-end by reconstructing the input images. Given an input image and its associated imaging hyperparameters (contrastive transfer function, and rotation and translation of the particle in the image estimated during *ab initio* reconstruction), CryoNeRF first reconstructs the 3D protein density and then projects this density according to the hyperparameters. CryoNeRF is then optimized by the mean-square error (MSE) loss by comparing the 2D projection of the predicted density and the 2D input image.

After training, CryoNeRF can be used to extract heterogeneous particle embeddings from all 2D particle images and predict the density for each protein particle in the dataset. In the case of heterogeneous datasets (**Fig. 1b**), the extracted particle embeddings can be further analyzed through UMAP embeddings to 2D, providing insights into the structural heterogeneity within the dataset.

### CryoNeRF for homogeneous reconstruction

We first benchmarked CryoNeRF’s ability to carry out consensus 3D density reconstruction on homogeneous datasets to testify the reliability of CryoNeRF reconstruction with widely used methods. We used the RAG1–RAG2 complex (EMPIAR-10049)^26^ and *Plasmodium falciparum* 80S ribosome (EMPIAR-10028)^27^, and the model was trained without the image encoder and the particle embedding. Following previous practices^20,29^, imaging hyperparameters were obtained from the datasets, and the particle poses were estimated by performing homogeneous refinement in cryoSPARC^19^. Here we compared with cryoDRGN^20^, a state-of-the-art deep learning-based Fourier-space cryo-EM reconstruction method, and cryoSPARC, a traditional voxel-based method.

Specifically, the model and the comparison methods were trained separately from two halves of each dataset to evaluate their reconstruction consistency and robustness. Following previous practice^30,31,^ two half-datasets were obtained by randomly splitting the 2D projection images into two sets. After training, we followed common practices^11,20^,32 to evaluate the resolution by calculating the gold-standard Fourier shell correlation (GSFSC) between the two reconstructions obtained from independent splits of the dataset. The GSFSC measures the agreement between the two maps across spatial frequencies. To determine the resolution, we used the standard GSFSC which measures the correlation between two densities at different spatial resolutions. The threshold was set to 0.143, and the resolution was identified as the point where the GSFSC curve dropped below this threshold. CryoNeRF achieved a higher resolution of 3.8 Å and 3.2 Å on EMPIAR-10049 and EMPIAR-10028, respectively. In comparison, cryoSPARC and cryoDRGN yielded resolution of 4.5Å / 4.0Å on EMPAIR-10049, and 4.1Å / 3.8Å on EMPAIR-10028, (see **Fig. 2a** and **d**). The related GSFSC curves are presented in **Fig. 2c**, where CryoNeRF yields consistently better performance compared to other methods. We further found, by comparing local resolution estimated from CryoNeRF and cryoSPARC reconstructions, that CryoNeRF achieves better local resolution in many regions (see **Fig. 2b**). These metrics clearly demonstrate CryoNeRF’s capacity for high-resolution reconstruction.

**Fig. 2.**
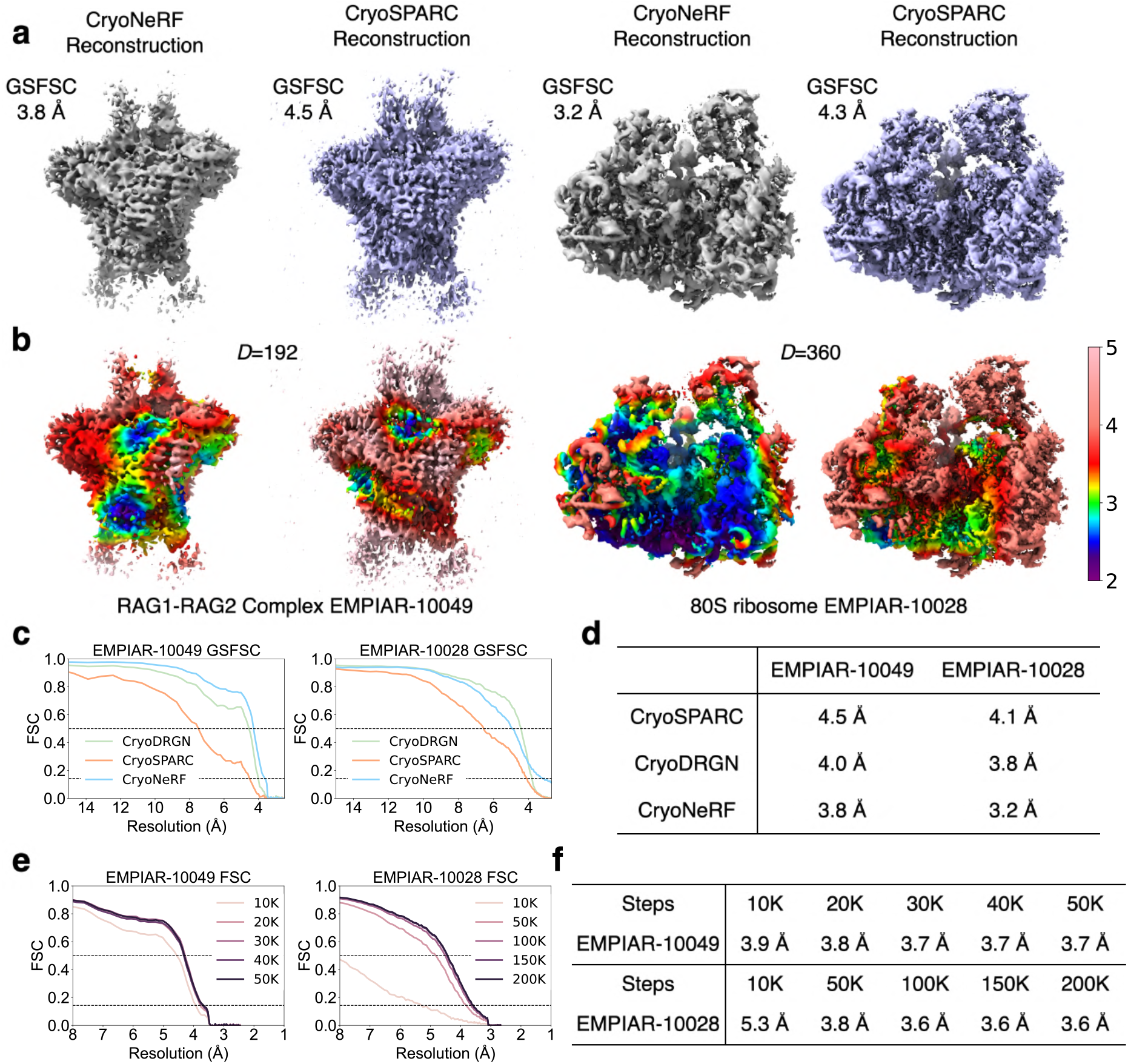
CryoNeRF reconstruction on homogeneous datasets. **a**, Density map comparison of the RAG1-RAG2 complex (EMPIAR-10049)^26^ and the *Pf*80S ribosome (EMPIAR-10028)^27^ reconstructed using CryoNeRF (grey) and the traditional voxel-based method cryoSPARC^19^ (purple). The gold-standard FSC (GSFSC) are computed from two half maps. **b**, Local resolution^28^ maps of CryoNeRF and cryoSPARC computed from half maps. **c**, GSFSC curve comparisons of different methods. **d**, Reconstruction resolution of different methods from GSFSC (FSC=0.143) **e**, The full map reconstruction Fourier shell correlation (FSC) curves of CryoNeRF compared with cryoSPARC reconstruction during training. **f**, Reconstruction resolution (FSC=0.143) of CryoNeRF in panel **e** relative to different training steps. Datapoints for FSC curves of different steps and GSFSC curves are in **Supplementary Table Tab EMPIAR-10049** and **Tab EMPIAR-10028**.

After validating the reconstruction quality of CryoNeRF, we further trained it with the full datasets using all available 2D images. Adopting the same setup as in previous work^20^, we used cryoSPARC as a reference method to evaluate our performance during training. We found that CryoNeRF converged well to a relatively high resolution (**Fig. 2e, 2f**) after iterative training, producing reconstructions correlated with the traditional methods at a resolution up to 3.7Å for EMPAIR-10049 and 3.6Å for EMPAIR-10028. This showcases the efficacy of using NeRF to represent and model the 3D structure for cryo-EM reconstruction.

### CryoNeRF for simulated conformational heterogeneity

We then evaluated CryoNeRF’s ability to resolve conformational heterogeneity in cryo-EM datasets by using the simulated IgG-1D dataset from CryoBench^32^. This dataset consists of 100,000 images from 100 atomic models with a continuous dihedral rotation of 360 degrees with an interval of 3.6 degrees, where 1,000 images are simulated for each rotation.

On the IgG-1D dataset, we trained the full CryoNeRF with the image encoder, which extracted particle embeddings from the particle images. After training, we embedded the extracted particle embeddings from images into 2D using UMAP^33^, and further colored the points with corresponding rotation angles (**Fig. 3a**). The embedded latent space forms a clear closed circle with smooth color variation between rotation angles, suggesting that CryoNeRF successfully resolved the conformational heterogeneity while preserving the spatial similarity between proteins in the latent space.

**Fig. 3.**
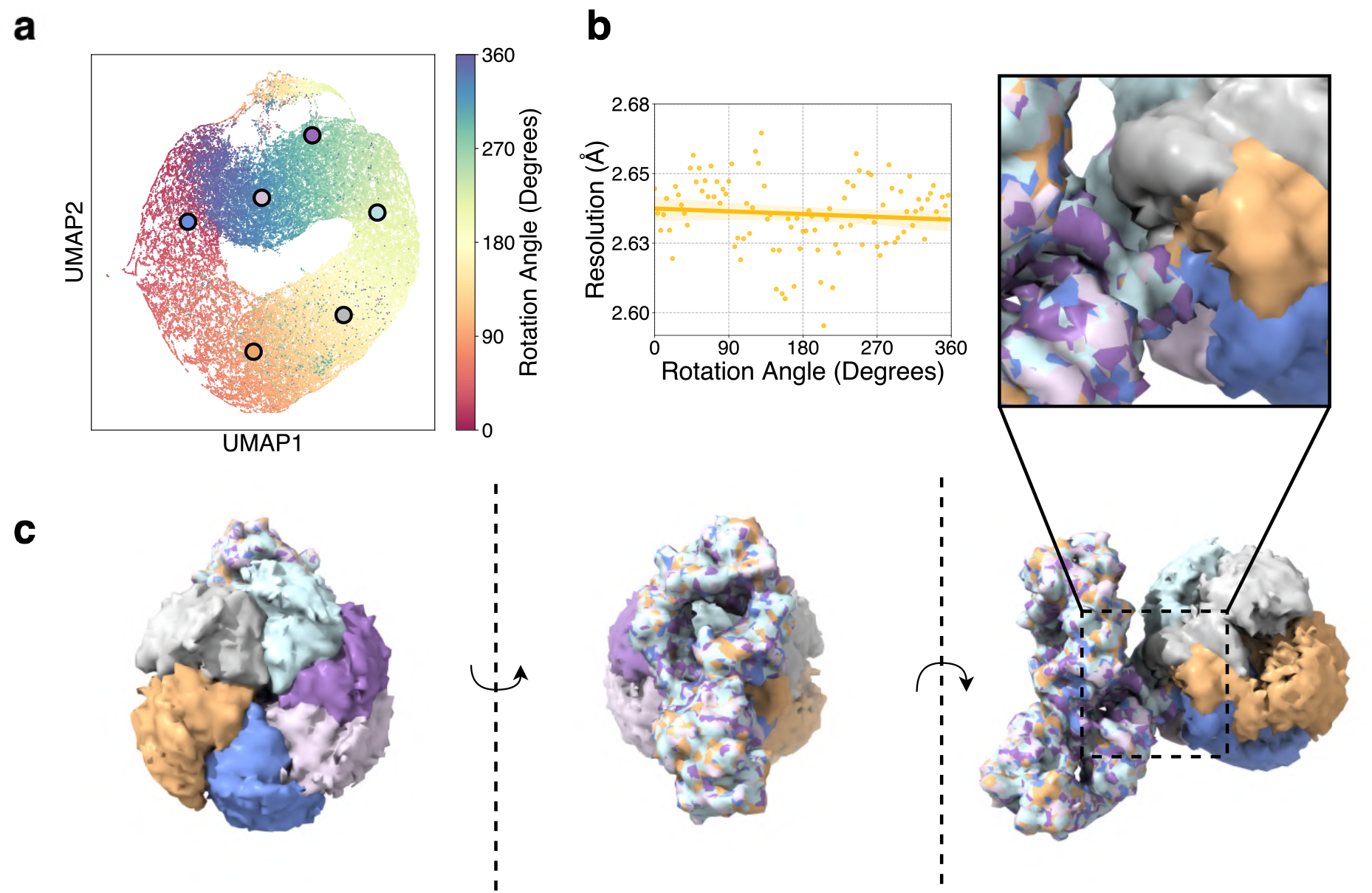
CryoNeRF on dataset with simulated conformational heterogeneity. **a**, UMAP visualization of particle embeddings on IgG-1D dataset, colored by the corresponding ground-truth rotation angle. **b**, The resolution (FSC=0.143) of CryoNeRF reconstructions of IgG-1D compared with groundtruths across different rotation angles. Here the reconstruction by CryoNeRF used the average particle embedding of each rotation angle. Each dots denote the performance of CryoNeRF at different angles. The solid line and shadow regions correspond to the regression line and the 95% confidence interval of regression, respectively. **c**, Composition reconstructions from sampled particle embeddings in the latent space, where the sampled regions correspond to the dots with same color. Additional density maps are shown in **Supplementary Video 1**. FSC datapoints for all rotation angles are in **Supplementary Table Tab IgG-1D**.

We further benchmarked the reconstruction resolution at different rotation angles. The comparison was done between the average density by CryoNeRF and ground-truth protein at each angle. The average protein density was obtained by taking the average of the corresponding particle embeddings at that angle. The results in **Fig. 3b** show that CryoNeRF achieves a relatively high resolution of better than 2.68 Å under all rotation angles. The average and variance of resolutions were 2.63 ± 0.01 Å, indicating that CryoNeRF achieved high-resolution and consistent reconstruction quality across all heterogeneous situations.

We then composited reconstructions (**Fig. 3c, Supplementary Video 1**) from sampled particle embeddings in **Fig. 3a**. The results on the left of **Fig. 3c** reveal that CryoNeRF effectively resolved the rotation heterogeneity and produced non-overlapping and evenly distributed reconstructions at the six sampled angles. We also observe that CryoNeRF successfully separated stationary and rotational regions, producing highly overlapping reconstructions at static areas (2^nd^/3^rd^ panel of **Fig.3c** and **Supplementary Video 1**). When investigating the boundary region between the stationary and rotational regions, a sharp boundary can be observed in the zoomed-in view.

In conclusion, CryoNeRF can clearly reconstruct conformational heterogeneity at a high resolution.

### CryoNeRF for simulated compositional heterogeneity

To test CryoNeRF’s ability to analyze complex compositional heterogeneity where particles come from different classes, we examined CryoNeRF on Ribosembly from CryoBench^32^, a dataset consisting of 335,240 images from 16 classes that correspond to a common core successively growing through the addition of proteins and ribosomal RNA^32,34^. In our experiments, CryoNeRF can generate heterogeneous particle embeddings by image encoder, aiming to capture the heterogeneity of particles in the dataset.

With the resulting trained model, we first generated 2D UMAP^33^ embeddings from the particle embeddings by CryoNeRF, colored by the corresponding ground-truth classes. We observed that different structural classes tend to cluster into different clusters, while similar structural classes tend to cluster together (**Fig. 4a**). We then averaged the embeddings from particle images of each class and reconstructed a protein density for that class from the averaged embedding. The CryoNeRF reconstructions of all classes (**Fig. 4d, Supplementary Video 2**) suggest that CryoNeRF successfully reconstructed different structures for different class, each of which exists significant structural differences relative to others, Meanwhile, CryoNeRF reconstructed similar structures for clusters that are close to each other in the UMAP embedding (**Fig. 4a**). The reconstructions also yielded high consistency with ground-truth densities^32^. The average resolution between reconstruction and ground-truth across different classes was 2.06 ± 0.08 Å, which was very close to the numerical limit of 2 Å resolution limit for Fourier shell correlation (FSC). The low variance and resolution of different classes in **Fig. 4b** indicate the stability of CryoNeRF on different classes. These results demonstrate the ability of CryoNeRF to produce high-resolution density maps by reconstructing them in 3D Euclidean space.

**Fig. 4.**
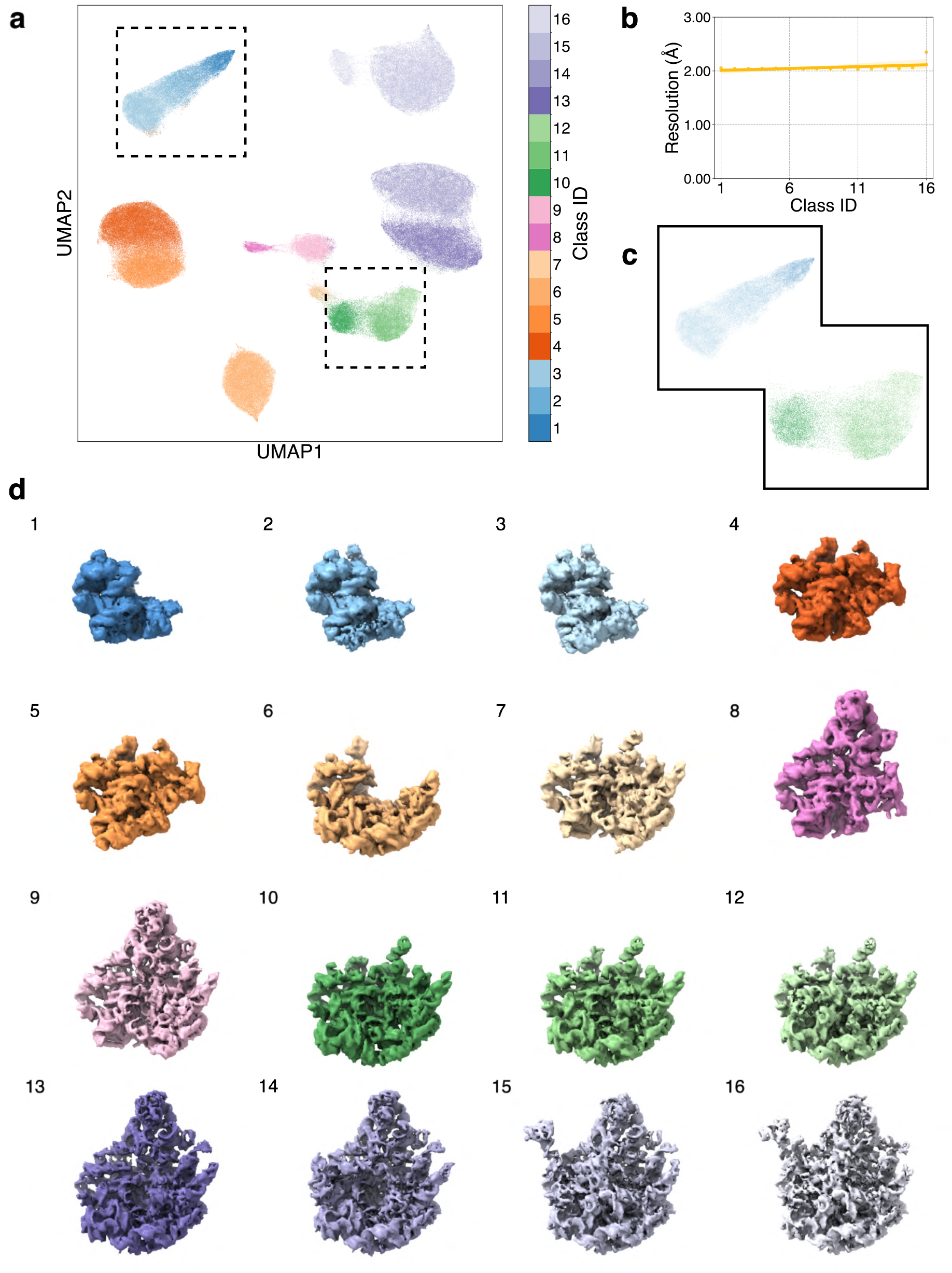
CryoNeRF on dataset with simulated compositional heterogeneity. **a**, Particle embeddings of Ribosembly embedded into 2D using UMAP. Figure is colored by the groundtruth class IDs. **b**, The resolution (FSC=0.143) of CryoNeRF reconstructions of Ribosembly compared with groundtruths over classes. CryoNeRF reconstructed using the average particle embedding of each class. Dots denote resolution at angles. The solid line and nearby regions are the regression curve of dots and the 95% confidence interval. **c**, Zoom in illustrations of regions marked on **a. d**, CryoNeRF reconstructed protein densities of all the classes. Proteins are colored with the same colors of colorbar in **a**. 360° videos of density maps are shown in **Supplementary Video 2**. FSC datapoints for all classes are in **Supplementary Table Tab Ribosembly**.

Notably, the learned latent space of CryoNeRF successfully captures the structural similarity across different classes. For example, as shown in **Fig. 4d** and **Supplementary Video 2**, Classes 1, 2, 3, and Classes 10, 11, 12 exhibit structures that are similar in overall shape but differ in specific regions, and this similarity was captured by CryoNeRF, mapping particle embeddings of these classes into close but separated positions (**Fig. 4a, 4c**). Through Euclidean 3D space reconstruction, CryoNeRF reflects the similarity of structures into learned particle embeddings.

### CryoNeRF for experimental conformational heterogeneity

We next tested CryoNeRF on a real-world experimental dataset to verify its capacity for resolving experimental conformational heterogeneity. Conformational heterogeneity is one of the most common types of heterogeneity and reflects the motion of the protein particle. This experiment used the pre-catalytic spliceosome (EMPIAR-10180)^23^, which includes 307,240 images.

We trained CryoNeRF with particle embeddings and visualized the resulting UMAP embeddings (**Fig. 5a**). In the UMAP embedding space, the particles form three main clusters, with the largest cluster showing an expected symmetry that is consistent with the process of undergoing conformational changes. To investigate the learned particle embeddings, we sampled particle embeddings corresponding to the cyan point and arrowhead/tail of arrows on the **Fig. 5a**, and reconstructed 3D conformations using the corresponding particle embeddings in **Fig. 5a**. In the corresponding **Fig. 5d, e, f**, clear movement can be observed, following the direction of arrows on **Fig. 5a**. The cyan point in **Fig. 5a** and the reconstruction in **Fig. 5b** are consistent with the previous work^20^ which identified a state in which the protein complex lacks the SF3b subcomplex. The grey arrow in **Fig. 5a** and reconstructions in **Fig. 5c** demonstrated a relatively small difference, implying that CryoNeRF clustered this region by identifying its stable spatial motion.

**Fig. 5.**
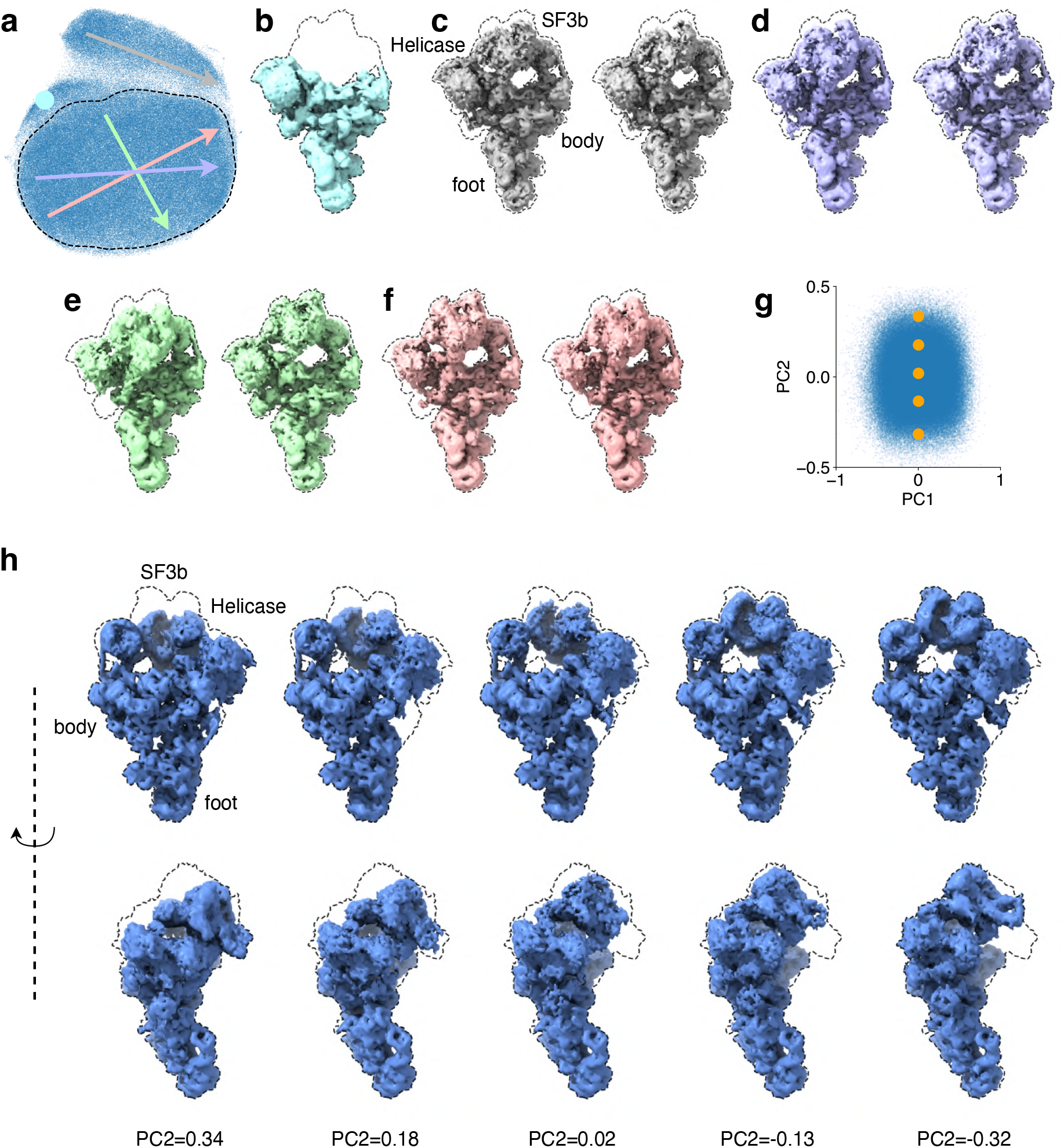
CryoNeRF on experimental conformational heterogeneous dataset. **a**, UMAP embeddings of particle embeddings from CryoNeRF. Points and ends of arrows show the sample position of the particle embeddings for reconstructions in **b-f**. The black dashed line represents the majority of protein particles, which is further analyzed in panel **g. b**, The reconstruction with the particle embedding at the cyan dot. **c-f**, Reconstructions with particle embeddings along the arrows with the corresponding colors. The left side of each panel shows the reconstruction from the embedding at the tail of the arrow, and the right side is from the embedding at the arrowhead. **g**, PCA embeddings of the dashed line region in **a. h**, Structures generated by traversing the selected region, as the orange dots marked in **g**. Additional density maps are shown in **Supplementary Video 3**.

To further investigate the latent space learned by CryoNeRF, we selected the largest cluster marked with the dashed line in **Fig. 5a** and embedded the corresponding particle embeddings to 2D with PCA (**Fig. 5g**) to extract the major movement pattern as the principal components. The PCA map shows a symmetric shape, consistent with the fact that this protein was in a conformational moving process. Then we traversed the 2D PCA embedding by sampling particle embeddings corresponding to points with varying PC2 values and the same PC1 value in the figure, and reconstructions viewed from front and side are shown in **Fig. 5h** and **Supplementary Video 3**. The reconstructions demonstrate clear continuous movement of the SF3b and Helicase regions.

In conclusion, the experiments on the EMPIAR-10180 dataset clearly showcased CryoNeRF’s ability to model and unravel the intricate conformational movements of proteins. By capturing subtle yet critical structural changes and projecting them into a well-organized latent space, CryoNeRF revealed dynamic transitions and movement patterns with striking clarity. The observed conformational heterogeneity, from large-scale motion to fine-grained structural variations, underscores CryoNeRF’s potential to push the boundaries of 3D reconstruction in structural biology.

### CryoNeRF for experimental compositional heterogeneity

Finally, we verified CryoNeRF’s ability to resolve experimental compositional heterogeneity. Compositional heterogeneity means that particles in a cryo-EM dataset exhibit distinct structures that can be grouped into different classes. This heterogeneity plays a crucial role in understanding molecular evolution and the functional divergence of biomolecules. In this experiment, we used Assembling LSU (EMPIAR-10076)^22^, which consists of 131,899 images from the bacterial large ribosome modular assembly process.

CryoNeRF was trained on the entire dataset with particle embeddings. UMAP was then applied to visualize the learned embeddings, which were colored according to the published major assembly states from the dataset^22^ (**Fig. 6a**). The UMAP embeddings formed six distinct clusters. In the EMPIAR-10076 study, six major states were identified (Class 1-6 in **Fig. 6a, c, Supplementary Video 4**), with a subset of particles unassigned to any class (represented as brown Class 0 in **Fig. 6a**). Due to the small size of Class 6 (1,853 particles) and its similarity to the unassigned Class 0 particles (26,575 particles), the UMAP embeddings of CryoNeRF for Class 6 overlapped with those of Class 0 (dashed line region in **Fig. 6a**). To further investigate the overlapping UMAP embeddings, we zoomed in on the overlapping region (**Fig. 6b)**. Here we can observe that CryoNeRF was able to cluster the Class 6 UMAP embeddings (blue) into a relatively condensed region, demonstrating its impressive ability to detect weak compositional heterogeneity signals.

**Fig. 6.**
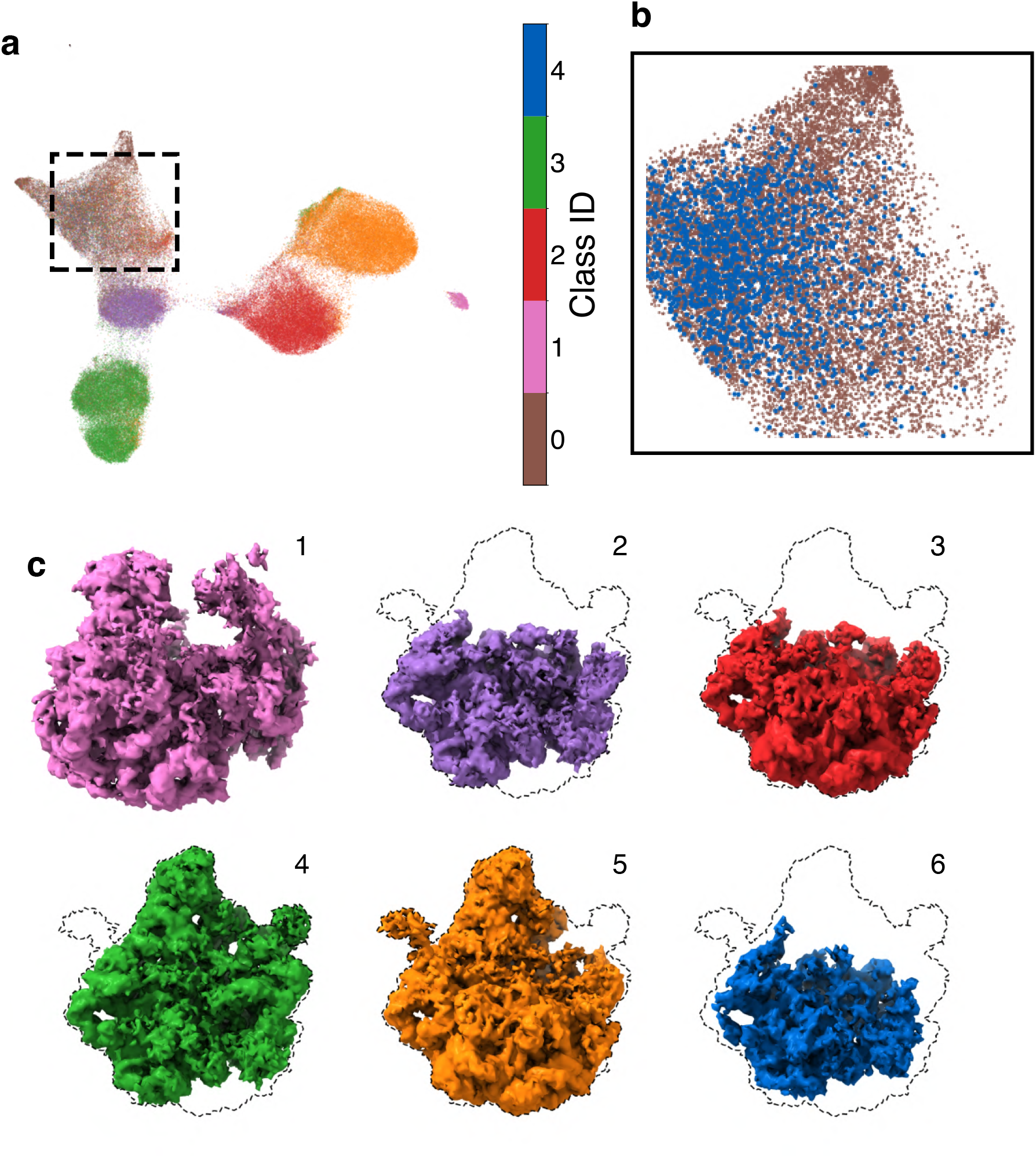
CryoNeRF on experimental compositional heterogeneous dataset. **a**, UMAP visualization of particle embeddings learned by CryoNeRF, colored according to the published major assembly states^22^. Particles without assigned classes are classified as Class 0 (brown), which overlap with Class 6 (blue) in the region outlined by the square dashed line. The dashed line region is enlarged for better visualization. **b**, The zoomed in view of the UMAP embeddings in the selected region in **a. c**, CryoNeRF reconstructions using the average particle embedding of each class. The density maps are color-coded to correspond with the class colors in panel **a**. 360° videos of density maps are shown in **Supplementary Video 4**.

Next, we generated reconstructions using the average particle embeddings learned by CryoNeRF for each class, with the resulting density maps shown in **Fig. 6c** and **Supplementary Video 4**. Notably, CryoNeRF successfully reconstructed the Class 1 70S ribosome, an impurity in the dataset consisting of only 2,018 particles (2% of the total dataset), directly from the full dataset without any filtering. In addition, all reconstructions in **Fig. 6c** and **Supplementary Video 4** recapitulate the known ribosome assembly process^22^.

In conclusion, CryoNeRF demonstrated remarkable performance in resolving experimental compositional heterogeneity in cryo-EM datasets. Using the LSU dataset (EMPIAR-10076), CryoNeRF identified six distinct assembly states through UMAP embeddings and was able to capture subtle heterogeneity signals. These results highlighted CryoNeRF’s power to disentangle and reconstruct complex heterogeneity directly from raw cryo-EM data.

## Discussion

CryoNeRF is a cryo-EM reconstruction method that operates directly in 3D Euclidean space, uniquely incorporating neural radiance fields (NeRF). Unlike recent neural network-based cryo-EM methods that rely on the Fourier slice theorem and perform reconstructions in Fourier space^20^, CryoNeRF avoids the numerical precision loss and high-frequency information degradation introduced by Fourier transformations. By reconstructing in Euclidean 3D space, CryoNeRF preserves the structural similarity in the learned particle embeddings, providing better interpretability. This work thus successfully bridges the gap between traditional cryo-EM methodologies and cutting-edge 3D computer vision research. Extensive experiments validated that CryoNeRF can produce high-resolution reconstructions on both homogeneous and heterogeneous datasets, effectively modeling conformational and compositional heterogeneity while maintaining interpretability through learned particle embeddings.

Despite its demonstrated effectiveness, the current version of CryoNeRF has limitations that require future improvements. Currently, CryoNeRF functions as a heterogeneity modeling tool^18,21,35,36^ within the traditional cryo-EM pipeline, relying on particle poses obtained from pre-performed homogeneous refinements. As a result, the reconstruction quality is influenced by the accuracy of the input poses. A promising future direction is to integrate pose estimation and search techniques to enable direct reconstruction from raw particle images. Additionally, CryoNeRF’s use of Euclidean 3D space reconstruction demands more GPU memory and computation compared to Fourier-based methods. For example, training on the IgG-1D dataset for 60 epochs took 18 hours on 4 A100 80GB GPUs. Addressing this challenge could involve developing more memory-efficient representations for NeRF.

We firmly believe that CryoNeRF will become a vital and accessible tool for cryo-EM reconstruction, serving as a reliable foundation for further macromolecular structure modeling.

## Acknowledgements

The authors thank Minkyu Jeon and Sukwon Yun for their discussion about datasets.

## Author contributions

XW and TC conceived the study, HQ and XW designed and implemented CryoNeRF and computed results. All the authors analyzed the results. HQ and XW drafted the manuscript and YZ, SW, WSN and TC edited it. All the authors read and approved the manuscript.

## Materials & Correspondence

Sheng Wang, William Stafford Noble, Tianlong Chen

## Code availability

The CryoNeRF source code is available at https://github.com/UNITES-Lab/CryoNeRF under GNU General Public License v3.0. CryoNeRF can run on Colab.

## Data availability

Trained CryoNeRF models, generated volumes and input files for training (excluding particle stacks) were deposited in Zenodo at https://doi.org/10.5281/zenodo.14602456. We used the following publicly available datasets: IgG-1D, Ribosembly, EMPIAR-10049 (Synaptic RAG1-RAG2 Complex), EMPIAR-10028 (*Plasmodium falciparum* 80S ribosome), EMPIAR-10076 (L17-depleted 50S ribosomal intermediates), EMPIAR-10180 (pre-catalytic spliceosome).

## Methods

### Cryo-EM reconstruction

Cryo-EM reconstruction aims to predict the 3D structures of protein particles in a dataset from 2D particle images of the dataset. In real experiments, particle images are collected as orthogonal projections of proteins illuminated by an electron beam.

The image formation model of a particle in cryo-EM can be described as

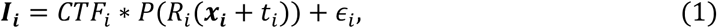

where *I*_*i*_ is the *i*-th image; *x*_*i*_ is the captured particle in that image (for simplicity, we refer to a selected region in the captured raw cryo-EM image that contains only one particle as a single “image”, as identified by the particle picking process^37-39^); *CTF*_*i*_ is the contrast transfer function, an experimental hyperparameter of the *i*-th image; *P* is the orthogonal projection operator; *R*_*i*_ is the rotation operator of the *i*-th particle relative to a pre-defined coordinate; and *t*_*i*_ is the translation of the *i*-th particle. The projected image is obtained with the following steps. First, the protein’s 3D density is translated by *t*_*i*_ and rotated by *R*_*i*_. Then the 3D density is projected to 2D space using the orthogonal projection operator *P*. The final image *I*_*i*_ is then available after being affected by *CTF*_*i*_ and combined with the inevitable noise *ϵ*_*i*_ produced during the imaging process in real-world experiments. For notational simplicity, we refer to this process as

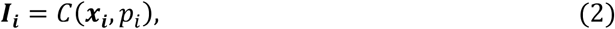

where *x*_*i*_ is the 3D density of the *i*-th particle and *p*_*i*_ = (*R*_*i*_, *t*_*i*_, *ϵ*_*i*_, *CTF*_*i*_) is the set of imaging hyperparameters.

The goal of cryo-EM reconstruction is to predict a 3D density 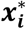 of the *i*-th particle that is close to the real map *x*_*i*_ by utilizing available 2D cryo-EM images 𝒥 = {*I*_*1*_, *I*_*2*_, …, *I*_*n*_}. Given an optimizable reconstruction function *𝒻*, cryo-EM reconstruction aims to minimize the difference between the reconstruction and the original map

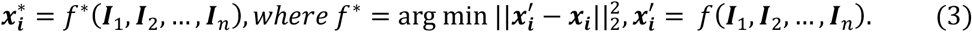

Here 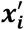 denotes a reconstructed density map of each particle, via the reconstruction function *𝒻, 𝒻*^*\**^ is the optimal reconstruction function, and 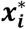 is the corresponding density map.

However, when doing reconstruction on images from real-world cryo-EM experiments, the real density map *x*_*i*_ is not available. Therefore, the problem is reformulated to optimize the reconstruction function by minimizing the error between 2D projections of the 3D reconstructed density and the available 2D projections 𝒥

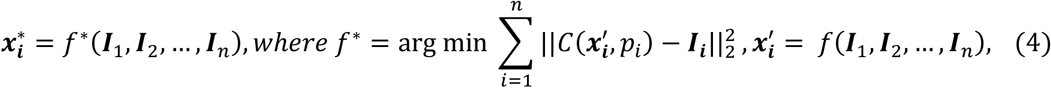

where 𝒥 = {*I*_1_, *I*_2_, …, *I*_*n*_} is the set of 2D particle images, and 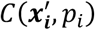 is the imaging process of the *i*-th particle. To further reduce the computational cost, instead of using images as the input to the reconstruction function, a grid of 3D coordinates *V* ∈ ℝ^*m*×3^ is employed to query density values, with *m* the number of 3D points determining the input. The optimization process directly optimizes density values on spatial points, namely

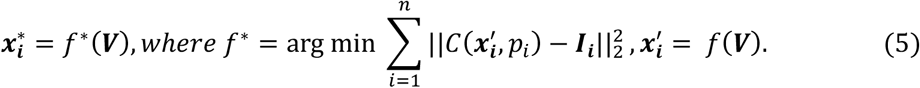

In our experiments, we set *m* = *h* × *h* × *h*, where *h* is the number of intervals on each edge of the grid volume.

### Cryo-EM reconstruction using NeRF

Following equation (5), in CryoNeRF we choose to use neural radiance field (NeRF) as the reconstruction function *𝒻*. NeRF is a function *𝒻*: ℝ^3^ ↦ ℝ that maps the 3D coordinate of a point into the density value of that point. Specifically, NeRF takes pose (*R*_*i*_ and *t*_*i*_ in cryo-EM reconstruction) as the input and predicts the projection of the 3D density at the given pose. The image formation process of NeRF is

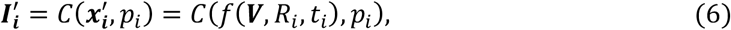

where 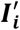 denotes the predicted image of the *i*-th particle, *R_i_* and *t_i_* are rotation and translation values corresponding to the particle in experimental image *I*_*i*_, and V is the coordinates of interest before being rotated and translated. NeRF first predicts density values on points of interest after rotation and translation, then generates an image by following the image formation model of cryo-EM.

However, in practice, equation (6) struggles to address the challenge of heterogeneity in cryo-EM reconstruction, because the grid V of 3D coordinates remains fixed and lacks the particle-specific information necessary for capturing heterogeneity within each particle *x*_*i*_. To overcome this limitation, we adopt the concept of the autoencoder—a neural network architecture specifically designed to learn latent embeddings from input data—and incorporate it into our NeRF-based reconstruction pipeline. This integration enables the extraction of structural heterogeneity information from input images. The image formation process of NeRF with latent embedding extraction is

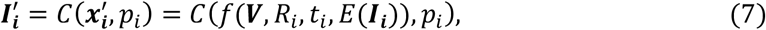

where *E*(*I*_*i*_) is the image encoder to extract heterogeneity information from *I*_*i*_. In CryoNeRF we implemented the image encoder *E*(*I*_*i*_) by using a standard ResNet-34^40^ architecture, with the last classification head removed, and a multi-layer perceptron (MLP) is added after the ResNet-34 to adjust the embedding dimension. After encoding, the embedding vector representing heterogeneity is concatenated with the queried density embedding from the multi-resolution hash encoding and then passed into the MLP decoder to predict the density value at the point (see **Fig. 1c**).

### Multi-resolution hash encoding-based neural representation

To boost the training speed and memory efficiency, multi-resolution hash encoding is used as the underlying neural representation for NeRF. Thus, the CryoNeRF method reconstructs a 3D map of a target protein by learning a multi-resolution hash encoding to represent the density distribution^25^.

A hash table is a data structure that stores key-value pairs. It uses a hash function *h*(⋅) to map a query into a key and uses the key to retrieve the value from the hash table. Hash tables are efficient, typically offering O(1) time complexity for searching and retrieving.

The multi-resolution hash encoding consists of hash tables arranged into *L* levels, and each level divides 3D space into small cubes called voxels, and voxels at different levels have different spatial sizes (see **Fig. 1d**). Each level *l* of the multi-resolution hash encoding contains *T*_l_ entries in the hash table, and each entry is an embedding vector with dimensionality *F*. The hash function *h*(⋅) in multi-resolution hash encoding takes the coordinates of a point as the input and maps it to an integer index as the key. Thereafter, the index can be used to retrieve the corresponding entries of the query point from levels of hash encoding.

Given a point ***p*** of interest, its coordinates are used as the query to the multi-resolution hash encoding to gather embedding vectors from this neural representation. At each level *l*, the voxel that contains point ***p*** is used to determine ***p***’s embedding at that level. Specifically, the hash function *h*(⋅) is applied to the eight vertices of the voxel in which the point ***p*** is located at level

*l*, mapping these vertices to embedding vectors stored in the hash table at that level. ***p*** ’s embedding is then calculated by doing interpolation of the embedding vectors from the eight vertices of the voxel it locates in (see **Fig. 1d**).

Embedding vectors of point ***p*** gathered from all the levels are concatenated, yielding an embedding vector for protein density of the point ***p*** with dimensionality *L * F*, and this vector is fed into a small MLP, along with the particle embedding extracted from the image encoder, to generate the final prediction of the density value of that point (see **Fig. 1d**).

### Implementation

The neural network of CryoNeRF was implemented with PyTorch^41^ and tiny-cuda-nn^42^. The multiresolution hash encoding was implemented with 16 levels of resolutions, and the embedding dimension was 2 on each level, resulting in a 32D density embedding. The image encoder used was a standard ResNet-34^40^, with the final classification removed for embedding extraction. Before feeding into the image encoder, a Hartley transform was first applied to the particle image. The extracted embedding from the image encoder was first fed to a 2-layer MLP with a hidden dimension of 128 to shrink the dimension, and the output particle embedding was 32D. The particle embedding was concatenated with all density embeddings and then input to the density decoder. The decoder was implemented as an MLP of 3 layers and 128 hidden neurons per layer. The model architecture was kept the same across all experiments for heterogeneity modeling. For the homogeneous reconstruction experiments in **Fig. 2**, the image encoder was removed, and the density decoder was decreased to 2 layers with a hidden dimension of 64. Training for all experiments were conducted with 4 NVIDIA A100 GPUs, and the training processing speed was 62.4 images/s for 128 * 128 particle images.

### Homogeneous reconstruction

In homogeneous reconstruction experiments, poses of particle images were pre-computed using cryoSPARC^19^ through *ab initio* reconstruction and homogeneous refinement tasks. To compute the GSFSC on EMPIAR-10049 or EMPIAR-10028, all particle images were randomly and evenly split into two non-overlapping parts, one reconstruction from each part of the dataset was produced, and GSFSC was computed as the FSC between two reconstructions from the two parts of the dataset. Thereafter, one reconstruction on each dataset was generated with the full dataset to compare with reconstructions produced by cryoSPARC^19^. All FSC curves were calculated without masking, and local resolution was estimated with Phenix^28^. The visualized protein densities were sharpened with the cryoSPARC sharpening tool, and the B-factor was estimated from two half maps.

### IgG-1D reconstruction

CryoNeRF was trained for 60 epochs on the full dataset of 100,000 images. The whole training process was finished in 27h. After training, the particle embeddings of all particle images were embedded into 2D using UMAP^33^, and colored with the ground-truth rotation angles from the dataset. After embedding, the 2D UMAP embeddings were divided into six clusters using K-means, and the six reconstructions produced with the centers of the clusters. Reconstructions for classes were produced by first averaging the embeddings of all particle images for a certain class, and then reconstructed with the averaged particle embedding, and finally compared with the ground-truth protein densities of each class.

### Ribosome reconstruction

The CryoNeRF architecture was also trained for 60 epochs on the full dataset with 335,240 images, which took around 90h. After training, the particle embeddings were projected into 2D with UMAP and colored with ground-truth classes. The average particle embedding of each class was used to reconstruct the protein density of that class. The colors of the 2D UMAP embeddings and protein densities followed the practice in CryoBench^32^, where a family of similar protein densities were colored with similar colors.

### Pre-catalytic spliceosome (EMPIAR-10180) reconstruction

We followed the same practice of heterogeneous reconstruction on simulation datasets. After training, we observed three clusters and first produced reconstructions from these clusters for exploration. We first examined the conformational movement of particles by traversing the cluster, but the movement was relatively small. We then extracted the particle embeddings from the largest cluster and re-embedded them with PCA, producing **Fig. 5g**. Given the symmetrical shape of the cluster in **Fig. 5g**, we traversed the cluster along its PC2 axis by sampling points, producing the protein densities in **Fig. 5h**.

### Assembling LSU (EMPIAR-10076) reconstruction

We trained CryoNeRF directly on the full dataset comprising 131,899 images. After training, the particle embeddings extracted by the image encoder were visualized using UMAP. The UMAP embeddings were colored based on the published major heterogeneity states for better interpretability.

